# Experience-dependent translational state defined by cell type-specific ribosome profiling

**DOI:** 10.1101/169425

**Authors:** Stephen M. Eacker, Khadijah Crawford, Lars Brichta, Markus Riessland, Nicholas T. Ingolia, Paul Greengard, Ted M. Dawson, Valina L. Dawson

**Affiliations:** Neuroregeneration and Stem Cell Programs, Institute for Cell Engineering, Johns Hopkins University School of Medicine, Baltimore, MD 21205, USA; Department of Neurology, Johns Hopkins University School of Medicine, Baltimore, MD 21205, USA; Diana Helis Henry Medical Research Foundation, New Orleans, LA 70130-2685, USA; Laboratory of Molecular and Cellular Neuroscience, The Rockefeller University, New York, NY; Department of Molecular and Cellular Biology University of California, Berkeley, Berkeley, CA; Department of Neuroscience, Johns Hopkins University School of Medicine, Baltimore, MD 21205, USA; Department of Pharmacology and Molecular Sciences, Johns Hopkins University School of Medicine, Baltimore, MD 21205, USA; Department of Physiology, Johns Hopkins University School of Medicine, Baltimore, MD 21205, USA

## Abstract

Experience-dependent neuronal activity regulates the translation of mRNA, supporting memory formation. We have developed a new method termed <u>t</u>ranslating ribosome affinity purification and <u>r</u>ibosome <u>p</u>rofiling (TRiP) which allows us to determine cell type-specific ribosome occupancy of mRNA with nucleotide resolution. Using TRiP we show that a memory-inducing experience creates a distinct translational state in mouse CA1 pyramidal cells. The experience-dependent translation state is characterized by enhanced translation of protein-coding open reading frames (ORFs) including numerous components of the actin cytoskeleton and calcium/calmodulin binding proteins, and by decreased translation of a defined subset of genes containing upstream ORFs (uORFs). Using animals heterozygous for an unphosphorylatable allele of the eukaryotic translation initiation factor 2α (eIF2α), we show that dephosphorylation of eIF2α contributes significantly to the experience-dependent translation state. These observations demonstrate that TRiP is a valuable methodology for studying physiologically relevant changes in translational state in genetically defined cell types.

## Introduction

Numerous studies over the last several decades have established the requirement for protein synthesis for the formation of new memories (Davis and Squire 1984). These observations have led to the widely accepted model that proteins synthesized in response to activity-dependent transcriptional and translational events are required for synaptic plasticity and long-term memory formation (Kelleher et al. 2004). Activity-dependent transcription can provide new mRNAs to be translated while changes in translation control which mRNAs are translated and how efficiently. Together both transcriptional and translational activation and/or suppression of specific mRNAs generates a ‘translational state’ that supports synaptic plasticity and memory formation.

Initiation of translation is the rate-limiting step in protein synthesis and the target of regulation by numerous signaling pathways (Sonenberg and Hinnebusch 2009). Translation initiation can be regulated both globally and in a transcript-specific manner shaping the overall translation state of a cell. Phosphorylation of eukaryotic translation initiation factor 2, alpha subunit (eIF2α) regulates both global translation initiation and translation of specific mRNAs (Ron and Walter 2007), and dynamic phosphorylation of eIF2α in the CNS plays an important role in behavior and synaptic plasticity (Costa-Mattioli et al. 2005; 2007; Di Prisco et al. 2014; Jiang et al. 2010; Huang et al. 2016). Notably, genetic reduction of eIF2α phosphorylation (Costa-Mattioli et al. 2007) or pharmacological reduction of p-eIF2α’s effects (Sidrauski et al. 2013) leads to improved performance in spatial and fear learning tasks. Increased levels of p-eIF2α are closely associated with memory impairment in rodent Alzheimer’s disease models, impairment that is reversed by genetic ablation of eIF2α kinases (Ma et al. 2013). Despite the clear importance of eIF2α phosphorylation in establishing a translational state that supports memory formation, there are few known p-eIF2α targets in the CNS.

Quantitative genome-wide assessment of the translational state has recently been made possible by the advent of ribosome profiling (Ingolia 2014). This method uses the recovery and sequencing of ribosome-protected RNA fragments to make a quantitative measurement of ribosome occupancy on mRNA at nucleotide resolution. However, defining the translational state associated with memory formation using ribosome profiling presents a number of technical challenges. Chief among them is the heterogeneity of cell types within any brain region, each cell type expressing a different collection of genes and receiving heterogeneous inputs in response to an experience. To reduce the problem of heterogeneity, we have utilized a cell type-specific ribosome immunopurification method called Translating Ribosome Affinity Purification (TRAP) (Doyle et al. 2008; Heiman et al. 2008). TRAP permits the purification of ribosome-mRNA complexes from genetically defined cell types with minimal manipulations that might affect the biochemical state of neurons. Taking advantage of TRAP and ribosome profiling methods, we have developed a new approach that allows us to define the translation state of genetically defined cell types within the mouse CNS. Using this method we show how a memory-inducing experience shapes the translational state of CA1 pyramidal neurons. Furthermore, we define how the phosphostatus of eIF2α contributes to the memory-associated translation state.

## Results

### TRiP defines cell type-specific translational state

The TRAP methodology enables the purification of translationally active mRNAs from a defined cell population, using affinity purification of enhanced green fluorescent protein-ribosomal protein L10a fusion (EGFPL10a) expressed transgenically under the control of a cell type-specific promoter. For these studies we selected the *Zdhhc2*-*EGFPL10a* transgene that is highly expressed in a subset of CA1 pyramidal cells of the hippocampus (Supplementary Figure 1A). mRNAs obtained by TRAP purification from the hippocampi from these animals show enrichment of EGFPL10a and significant depletion of inhibitory neuron and glial transcripts (Supplementary Figure 1B). TRAP-purified polyribosomes were subjected to RNase I digestion and ribosome footprints were recovered and sequenced (Figure 1A). Ribosome footprints demonstrated enrichment in coding regions and a high degree of 3 nt periodicity (Supplementary Figure 1C) and technical replicates from independent experiments exhibited substantial reproducibility (R = 0.9887, Figure 1B). We performed ribosome profiling from total hippocampal lysates and compared to the TRAP-purified profile (Fig 1C). Among genes most enriched by TRAP was *Zdhhc2*, the gene driving EGFPL10a expression. Genes depleted from TRAP samples include markers of glial cell types (*Gfap*, *Aldh1l1*, *Olig1*, *Mag*) and inhibitory neurons (*Gad1*, *Gad2*, *Sst*, Supplementary Table 1). These data indicate successful recovery of ribosome footprints from the excitatory pyramidal cells of the CA1. We term this new methodology <u>T</u>RAP <u>R</u>ibosome <u>P</u>rofiling (TRiP).

**Figure 1:**
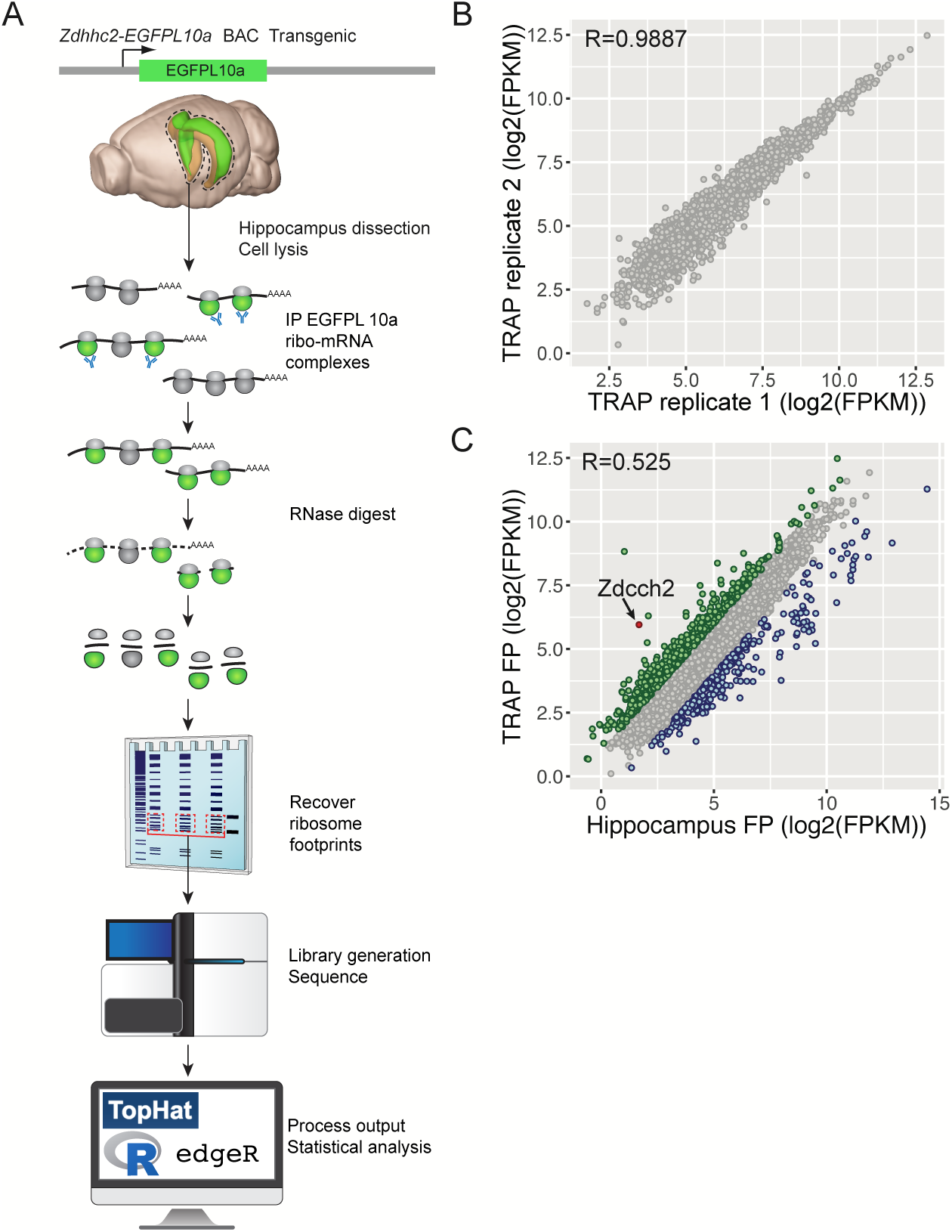
Translating Ribosome Affinity Purification and Ribosome Profiling (TRiP). (A) Schematic workflow of TRiP. (B) Comparison of replicates of TRiP showing Pearson’s correlation (R value). Replicates were performed on separate days to assess reproducibility and reveal any potential batch effects. (C) Comparison of results from TRiP from *Zdhhc2*-*EGFPLl0a* BAC transgenics and total hippocampus ribosome profiling. mRNAs enriched in TRiP samples greater than 2-fold are shown in green while those in blue represent greater than 2-fold depletion relative to total hippocampus profile. Pearson’s correlation (R value) displayed for this comparison.

### Identification of translated ORFs

We took advantage of codon-level resolution that TRiP affords to annotate translated open reading frames (ORFs) in *Zdhhc2*^+^ CA1 neurons using Ribotaper (Calviello et al. 2015). This software borrows the multitaper algorithm from signal detection theory to identify statistically significant periodicity in ribosome profiling data to identify actively translated segments of RNA. We applied Ribotaper to TRiP data, limiting our analysis to well-phased ribosome footprints 28-30 nt in length (Supplementary Figure 2A). Of ~55M unique TRiP footprints, we identified ~20M unique predicted P-sites (Supplementary Figure 2B). The TRiP footprints show clear enrichment of in-frame P-sites in consensus coding sequences (CCDS) compared to UTRs and non-coding RNAs (Supplementary Figure 2C), with overall 91% agreement with annotated CCDS definition (Supplementary Figure 2D).

Ribotaper identified 17,730 statistically significant ORFs within 12,904 genes, an average of 1.37 ORFs per gene (*p* < 0.05, Supplementary table 2). The majority of the ORFs identified by Ribotaper correspond to the canonical ORF annotated for a gene (56.6%, Fig 2A). The next most common class of ORFs identified are ORFs initiated from internal N-terminal methionine codons (niORFs, 35.4%, Fig 2B). While there are likely cases of internal initiation events, the majority of niORFs are likely the result of alternative 5’ ends or low coverage of 5’ ends of the canonical ORF. There are also a small number (0.6%) of ORFs that exhibit an N-terminal extension of the canonical ORF (neORF, Fig 2b) which may be the result of alternative 5’ ends or incorrectly annotated translation initiation sites. Ribotaper also identified upstream ORFs (uORFs) which account for 6% of all ORFs identified and includes previously identified uORFs such as observed in the NMDA receptor 2a mRNA (*Grin2a*, Fig 2B, (Wood et al. 1996). The remaining class of translated ORFs lie downstream of the canonical (dORFs, 1.4%), which in some cases are robustly translated (Fig 2B). Because sequences corresponding to niORF and neORF translation overlap nearly completely with the canonical ORF and are therefore indistinguishable, we have combined results for these three groups and refer to them as the main protein coding ORF (mORF) for any given transcript.

**Figure 2:**
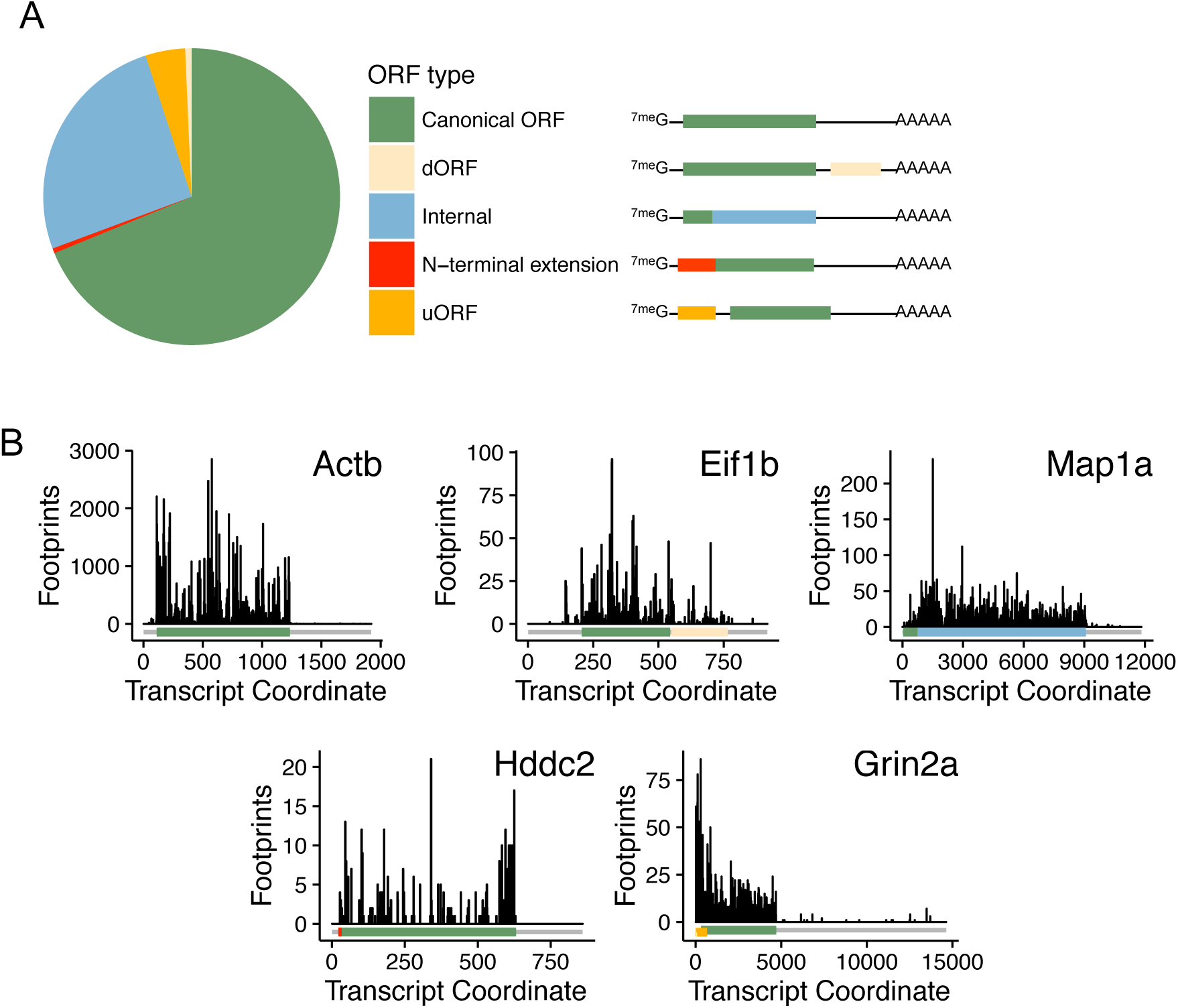
Ribotaper annotates translated open reading frames in Zdhhc2+ CA1 neurons. (A) Proportion of open reading frame (ORF) classes identified by Ribotaper. ORF classes include the canonical annotated ORF, ORFs that initiate downstream of annotated initiation sites (Internal) or upstream of annotated initiation sites (N-terminal extension). ORFs that do not share reading frame with the canonical ORF are referred to as either upstream ORFs (uORFs) or downstream ORFs (dORFs). (B) Example gene models of each class of identified ORF.

### Defining translational state associated with memory formation

Using Ribotaper annotations, we sought to define how a memory-inducing experience altered the translational state of neurons. We selected the well-characterized fear conditioning paradigm as a physiologically relevant task to measure how memory-inducing stimuli alter the translation state of hippocampal CA1 neurons. Fear conditioning memory requires protein synthesis for the formation of long-term but not short-term memory (Bourtchouladze et al. 1998; Schafe et al. 1999). Following habituation to the context, *Zdhhc2*-*EGFPL10a* transgenic mice were exposed to a 30 sec tone followed by a mild foot shock and then returned to their home cage (Figure 3A). This paradigm induced robust long-term memory as determined by tone-induced freezing 24h after training (Supplementary Figure 3). To determine how the fear conditioning paradigm alters the translational state of CA1 neurons, we performed TRiP on hippocampus 1h following the fear training period and compared to naïve mice. This time point was selected because it is at the transition between early-LTP and protein synthesis-dependent late-LTP and the time at which experience-dependent bulk protein synthesis peaks (Matthies 1989). A total of 421 ORFs showed significant increase in translation and 199 ORFs showed a decrease (Figure 3B, >1.5-fold, FDR <0.05, *p* < 0.01, Supplementary Table 3).

**Figure 3:**
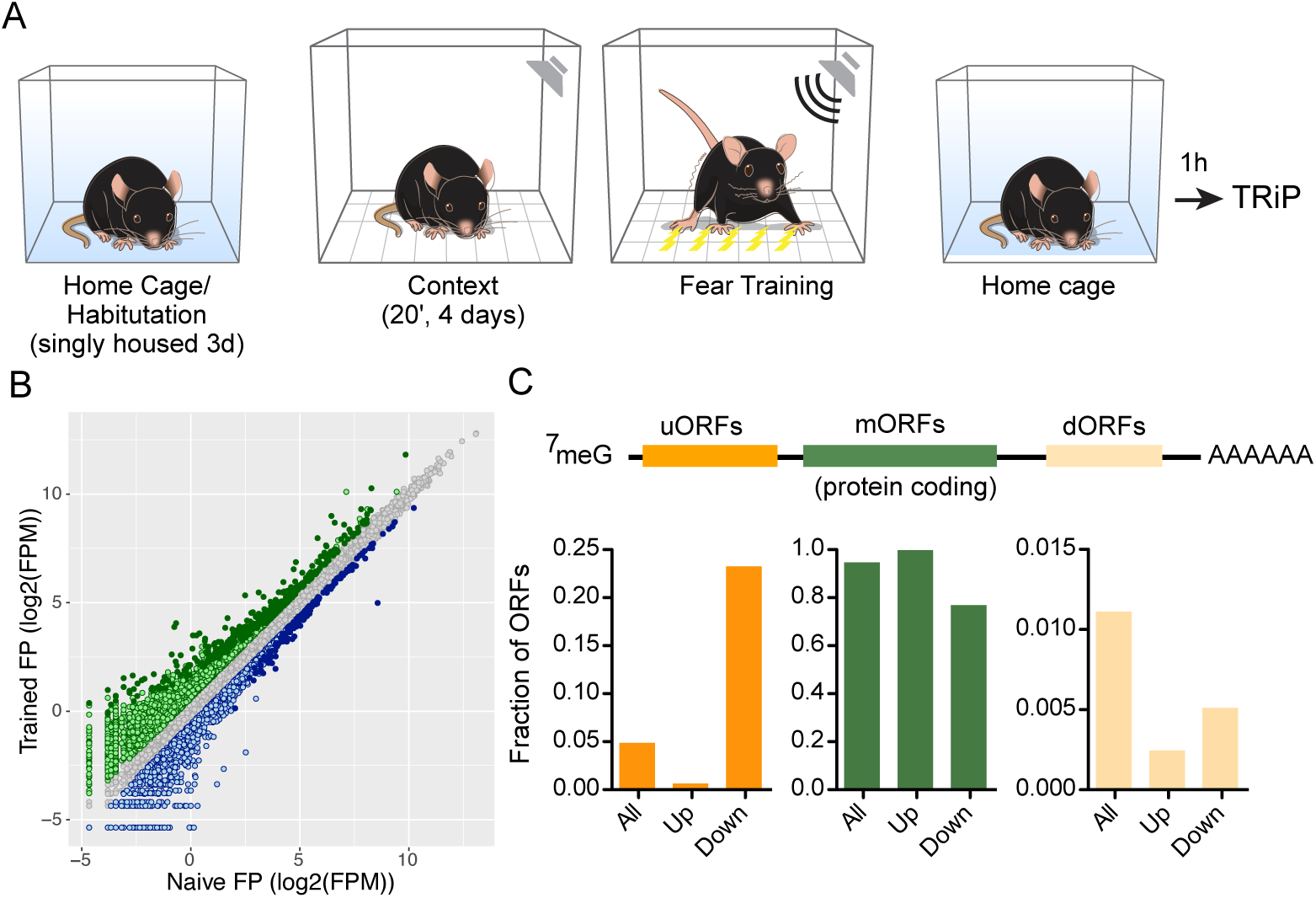
TRiP determines the translation state of CA1 pyramidal cells following a memory-inducing experience. (A) A schematic describing the behavioral paradigm used to generate a memory-induced translation state. (B) Comparison of differentially translated ORFs following fear training. Each point represents the mean footprints per million (FPM) value for trained (n=3) or naïve (n=3) mice TRiP experiments. Light green points reflect >1.5 fold increased translation of an ORF while dark green are ORFs show statistically significant increase in translation (FDR<0.05, *p* <0.01, edgeR GLM testing). Conversely, light blue points represent ORFs showing >1.5 fold decreased translation and dark blue show statistically significant decrease in translation (FDR<0.05, *p* <0.01, edgeR GLM testing). (C) Proportion of ORF classes showing statistically significant decrease (Down) or increase (Up) in translation following fear training. ORF classes showing increased and decreased translation differ significantly than expected based on the proportion of classes of all ORFs identified (^∗∗∗^ *p* < 0.001, Fisher’s exact test, see text for details).

Among the mRNAs demonstrating enhanced translation, gene ontology analysis (Supplementary Figure 4) identified an enrichment of genes involved in the cytoskeleton protein and actin binding (corrected *p* < 1.2×10^−6^ and 8.3 ×10^−6^ respectively) consistent with changes in structural plasticity associated with establishment of long-term memory (Rudy 2015). There was also an observed enrichment in calcium and calmodulin binding proteins (corrected *p* < 0.0077 and < 0.037) also consistent with the essential role of calcium/calmodulin signaling during synaptic plasticity. Among the downregulated ORFs there was an enrichment of DNA binding and transcription regulatory activity proteins (corrected *p* < 9×10^−7^ and 8.8×10^−3^ respectively), which included the immediate-early transcription factors Fos, Egr1, Egr4, Junb, and Npas4. Though seemingly contradictory to their known role in synaptic plasticity, this phenomenon has been previously observed (Cho et al. 2015) and may represent a reset in the transcriptional landscape following a burst in activity-dependent transcription (see Discussion).

Classes of ORFs differentially translated following training showed significant biases (Figure 3C). Both uORFs and dORFs were substantially depleted from ORFs that were significantly upregulated following training (10-fold and 4.7-fold respectively, *p* < 0.001, Fisher’s exact test). Conversely, uORFs are enriched among ORFs showing significant decrease in translation following training (4.8-fold, *p* < 0.001 Fisher’s exact test). The presence of uORFs has previously been shown to modulate the translation of downstream protein-coding mORFs, although this is not a universal feature of uORF-mORF interaction. Overall we detect no statistically significant effect on the translation of mORFs of genes containing uORFs following training compared to non-uORF containing genes (Supplementary Figure 5A). Interestingly, we find that dORF-containing genes show a significant reduction in mORF translation following training (Supplementary Figure 5B, *p* < 6.4 × 10^−9^, Wilcoxon rank sum tests). In summary, TRiP analysis shows that a memory-inducing experience alters the translational state of CA1 pyramidal cells, increasing translation of mRNA involved in structural plasticity and calcium/calmodulin signaling and depressing translation of some immediate-early transcription factors. The memory-inducing experience also selectively impairs uORF and dORF translation while promoting utilization of protein-coding mORFs.

### eIF2α dephosphorylation drives the pro-memory translational state

Changes in translational state following training are known to be the result of the integration of numerous signaling pathways that modulate both transcriptional (West et al. 2002) and translation control (Costa-Mattioli and Sonenberg 2008). However, selective translation of non-canonical ORFs in higher eukaryotes is a poorly understood phenomenon. One relative well-studied mechanism of differential translation of ORFs is mediated by the phosphorylation of the eukaryotic initiation factor 2, alpha subunit (eIF2α). eIF2α is a component of the eIF2 complex, that when associated with the initiator methionine tRNA and GTP forms a ternary complex required for translation initiation (Hinnebusch et al. 2016). The ternary complex promotes translation initiation through its participation in the 43S pre-initiation complex (PIC), which scans the 5’ end of mRNA and binds the start codon. When eIF2α is phosphorylated, the affinity of eIF2 for its GEF (eIF2B) is dramatically increased, sequestering the GEF and preventing regeneration of the mature ternary complex. In some cases, uORF translation impairs the translation of downstream mORFs. However, under conditions of high eIF2α phosphorylation, the 40S ribosomal subunit is thought to scan through the uORF, thus permitting the initiation of a mORF. Thus, eIF2α phosphorylation sculpts the translational state of cells by regulating global translation initiation and selective translation of ORFs.

We confirmed previous reports that training animals in a fear conditioning task results in decreased levels of phosphorylated eIF2α (p-eIF2α), reducing p-eIF2α levels to <50% of naïve levels (Figure 4A) (Costa-Mattioli et al. 2007). This training leads to p-eIF2α levels comparable to those observed in animals heterozygous for an unphosphorylatable allele of eIF2α (*eIF2s1*^*S51A*/+^). To probe the role of eIF2α phosphorylation in determining the translation state of CA1 pyramidal cells, we performed TRiP on naïve *eIF2s1*^*S51A*/+^ reasoning that the translational state of these animals would allow us to determine the effect of reduced p-eIF2α in the absence of other activity-dependent processes.

**Figure 4:**
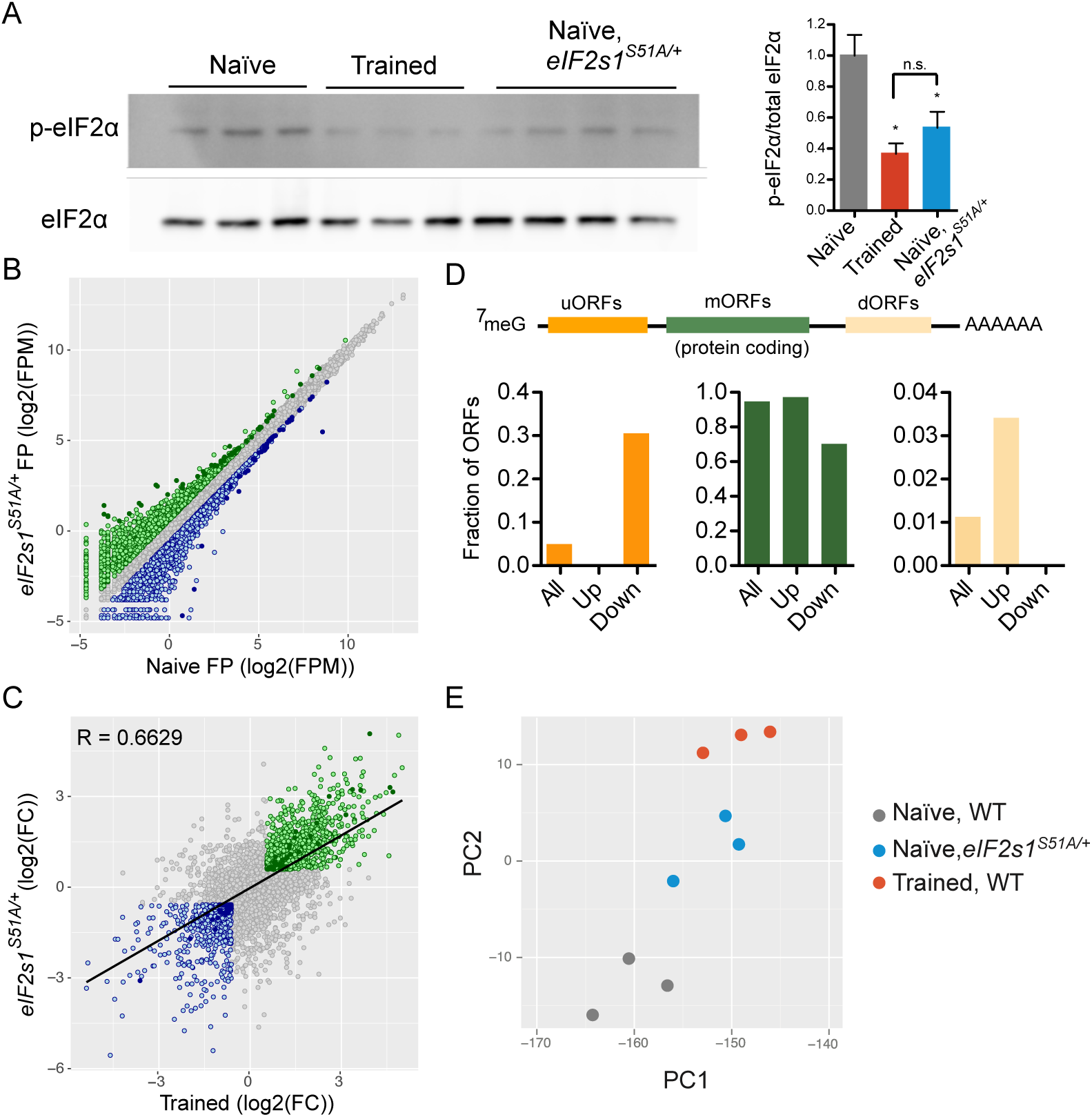
eIF2α phosphostatus partially recapitulates the training effect on translation state. (A) Training reduces levels of p-eIF2α levels in the hippocampus. Mice heterozygous for an unphosphorylatable mutant eIF2α (*eIF2s1S51A*/+) show levels of p-eIF2α similar to mice after training (^∗^ *p* < 0.05 compared with naïve wild type, one-way ANOVA, with Boneferoni post-test, n = 3-4). (B) Comparison of differentially translated ORFs between naïve wild-type of *eIF2s1S51A*/+ mice. Each point represents the mean footprints per million (FPM) value for *eIF2s1S51A*/+ (n=3) or naïve (n=3) mice TRiP experiments. Light green points reflect >1.5 fold increased translation of an ORF while dark green are ORFs show statistically significant increase in translation (FDR <0.05, *p* <0.01, edgeR GLM testing). Light blue points represent ORFs showing >1.5 fold decreased translation and dark blue show statistically significant decrease in translation (FDR <0.05, *p* <0.01, edgeR GLM testing). (C) Comparison of the fold-change (FC) in translation comparing effects *eIF2s1S51A* mutation and training. Color coding is a described in (B) but applies only to ORFs showing changes in both *eIF2s1S51A*/+ and trained groups. Line is a linear regression and reporting Pearson’s correlation (R). (D) Proportion of ORFs showing significant changes in translation in *eIF2s1S51A*/+ mice. Proportions of ORF classes showing increased and decreased translation differed significantly from expectations (^∗∗∗^ *p* < 0.001, Fisher’s exact test, see text for details). (E) Plot of principal components (PC) describing primary source of variation between TRiP samples from wild-type (WT) naïve, wild-type trained, and *eIF2s1S51A*/+ mice.

In total we identified 59 ORFs that demonstrated significant increase in translation and 33 that showed significant decrease in translation in *eIF2s1*^*S51A*/+^ mice compared to naïve wild-type mice (Figure 4B, Supplementary Table 3). Overall, the changes in translation following training and resulting from the *eIF2s1^S51A^* mutation are strikingly similar (Figure 4C-E). There is a significant overlap in significantly differentially translated ORFs between *eIF2s1*^*S51A*/+^ and trained wild-type mice (62/94 overlap, *p* < 6.1×10^−7^, Fisher’s exact test). ORFs with increased translation in *eIF2s1*^*S51A*/+^ show the greatest overlap with trained mice (46/59, *p* < 1×10^−6^, Fisher’s exact test), while ORFs with decreased translation showed only statistically modest, but still a majority overlap (20/33, *p* = 0.052, Fisher’s exact test). Of the genes showing decreased translation in *eIF2s1*^*S51A*/+^ mice, 27% contain at least one uORF detected by Ribotaper, as compared to 9% of all genes expressed. This suggests that at least in some cases, regulation of translation is occurring via the canonical uORF-mediated mechanisms ascribed to p-eIF2α (Hinnebusch et al. 2016). As was observed following training, there is an enrichment of uORFs among the ORFs showing significant decrease in translation in *eIF2s1*^*S51A*/+^ mice (6.8 fold, *P* < 0.001, Fisher’s exact test). No uORF appears among ORFs showing increased translation in *eIF2s1*^*S51A*/+^ mice, further suggesting that eIF2α phosphostatus significantly impacts ORF selection during translation initiation.

As observed following training, TRiP analysis of *eIF2s1*^*S51A*/+^ mice suggests that non-canonical ORF translation may influence mORF translation. Translation of mORFs in mRNAs that contain a uORF are slightly decreased in *eIF2s1*^*S51A*/+^ (Supplementary Figure 6A, *P* < 0.05, Wilcoxon rank sum test), a phenomenon not observed following training. However, as was observed following training, a more pronounced reduction in mORF translation is observed in dORF-containing mRNAs (Supplementary Figure 6B, *P* < 6.4 ×10^−9^, Wilcoxon rank sum test).

To compare the overall translational state of CA1 cells between naïve, trained, and *eIF2s1*^*S51A*/+^ mice, we performed principal component analysis (PCA) on the TRiP results. We represent the majority of the variance between these experimental groups in a two-dimensional PCA vector plot. We observed that individual samples are clustered closely by group with the most differences observed between naïve and trained samples. The PCA shows that the translational state of *eIF2s1*^*S51A*/+^ CA1 neurons sits in an intermediate position (Figure 4E). Similar results were observed when the data were analyzed using multi-dimensional scaling, another dimension reduction approach (Supplementary Figure 6). These data suggest that the phosphostatus of eIF2α is a significant contributor to the translational state of CA1 following a memory-inducing experience.

### Targets of eIF2α-mediated repression

Phosphorylation of eIF2α regulates translation initiation by two known mechanisms. As described above, eIF2α plays a role in ORF selection during translation initiation. However, eIF2α phosphorylation principally regulates global translation initiation of mORFs by modulating the availability of 43S PIC. Consistent with eIF2α’s role in general translation initiation, we observed that global protein synthesis is elevated in the hippocampus of *eIF2s1*^*S51A*/+^ compared to wild-type littermates (Figure 5A). To investigate how eIF2α phosphorylation can influence transcript-specific translation control, we generated mouse embryonic fibroblasts (MEFs) from *eIF2s1^S51A/S51A^* (A/A) and *eIF2s1*^+/+^ (S/S) littermates. As observed in the hippocampus, genetic impairment of eIF2α phosphorylation leads to a significant increase in general translation (Figure 5B).

**Figure 5:**
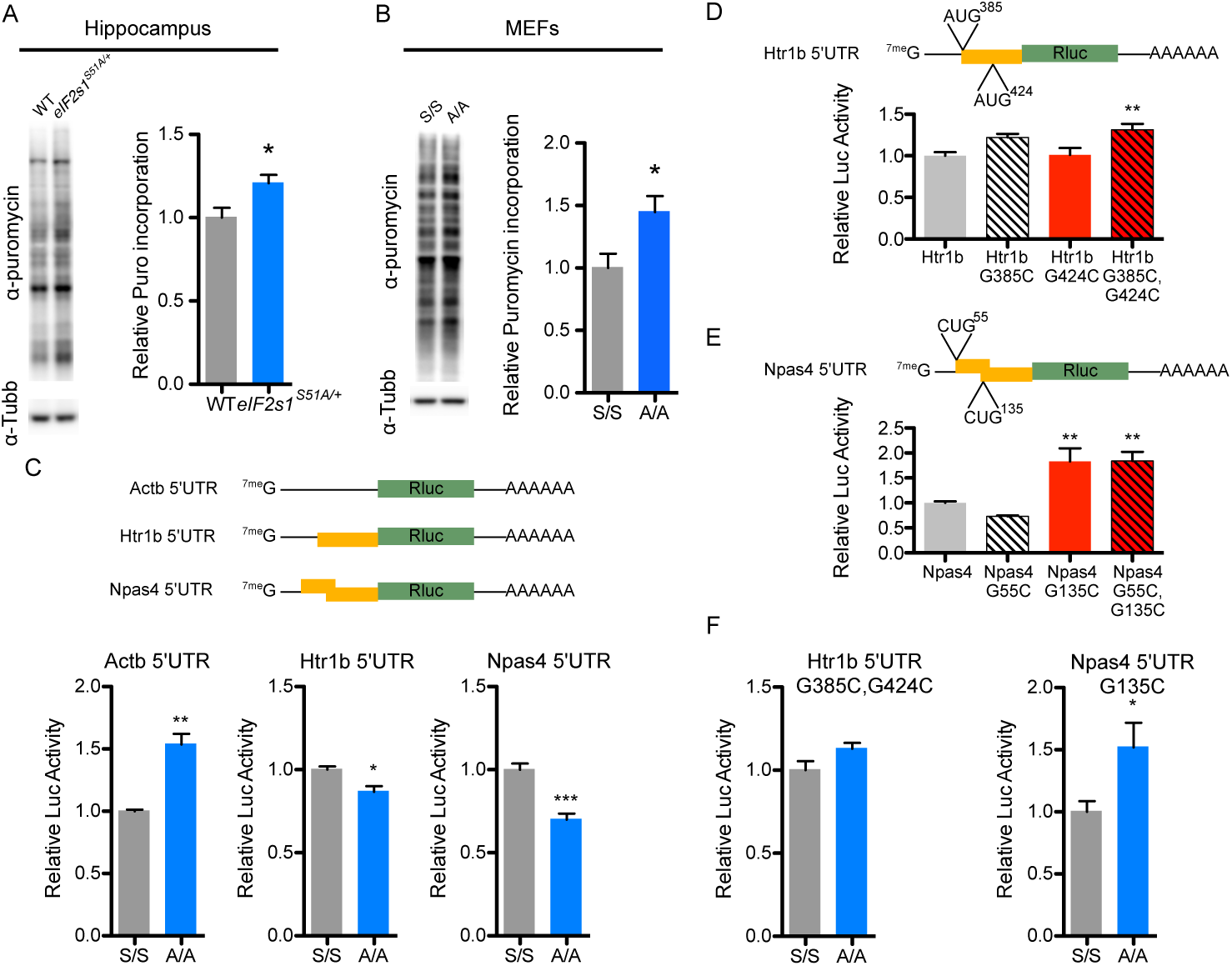
eIF2α phosphorylation regulates global and transcript-specific translation. (A) Puromycin-release assay from hippocampal lysates shows increased protein synthesis in *eIF2s1S51A*/+ (^∗^ *p* < 0.05, unpaired T-test, n = 7 per group). (B) SunSET assay in mouse embryonic fibroblasts (MEF) wild-type (S/S) or homozygous for *eIF2s1S51A* allele (A/A, ^∗^ *p* < 0.05, unpaired T-test, n = 5 per group). (C) Schematic of 5’UTR Rluc reporter constructs with orange blocks representing uORFs. Luciferase assays show that uORF-containing 5’UTR reporters show decreased translation in A/A MEFs (^∗^ *p* <0.05, ^∗∗^ *p* <0.01, ^∗∗∗^ *p* <0.001, unpaired T-test, n= 6 per group). (D) Mutational analysis of Htr1 b 5’UTR (^∗∗^ *P* <0.0l, one-way ANOVA with Boneferroni post-test, n = 9/group). (E) Mutational analysis of Npas4 5’UTR (^∗∗^ *p* <0.01, one-way ANOVA with Boneferroni post-test, n = 9 per group). (F) Mutations of uORF start codon eliminates repressive p-eIF2α-dependent effect of Htr1 b and Npas4 5’UTR reporters (^∗^ *p* <0.05, unpaired T-test, n = 9 per group).

We reasoned that mRNAs containing uORFs that were significantly repressed in both trained and naïve *eIF2s1*^*S51A*/+^ mice would be potential physiologically relevant candidates for this mechanism of regulation. To test this hypothesis, we generated a series of *Renilla* luciferase (Rluc) reporters bearing the 5’ UTRs of selected targets. When transfected into A/A MEFs, the Rluc reporter containing the *beta*-*actin* (*Actb*) 5’ UTR, which has no uORF, is translationally upregulated, consistent with the associated increase in general translation (Figure 5C). By contrast, reporters for two uORF containing 5’UTRs, *serotonin receptor 1B* (*Htr1b*) and *neuronal PAS domain protein 4* (*Npas4*), show significant reduction in translation in A/A MEFs compared to S/S (Figure 5D and E). Htr1b contains two uORFs in frame with each other identified by Ribotaper initiating from an AUG at 383 nt and 422 nt respectively (Supplementary Figure 8). To determine if these uORFs play a role in the repression of eIF2α-dependent translation, we mutated the G at 385 and 424 to C (G385C and G424C) individually and together. We found that the G385C mutation lead to modest but insignificant increase in reporter translation while G424C had no effect. However, when combined with G385C, G424C showed a statistically significant increase in reporter translation (Figure 5F). Similarly, Npas4 contains two potential uORF sequences that initiated from non-canonical CUG initiating sites (Supplementary Figure 7). Mutating the first potential initiation site (G55C) had no effect on reporter translation while mutating the second CUG (G135C) robustly increased reporter translation (Figure 5G). Combining both Npas4 5’UTR mutations had no further effect than the G135C mutation alone. The Htr1b G385C, G424C and Npas4 G135C mutant reporters eliminated the repressive effect observed in A/A MEFs suggesting that eIF2α regulates Htr1b and Npas4 through a uORF-mediated mechanism.

## Discussion

Protein synthesis is essential for the establishment of long-term memory. Other than a handful of immediate-early genes (IEGs) that are transcriptionally induced by neuronal activity, relatively little is known about what proteins are synthesized during the formation of new memories. This is in part due to the complex milieu of cells in the brain, each of which having a potentially different translational response to experience-induced neuronal activity. In this study we introduce a new methodology, TRiP, to determine the translation state of genetically defined populations of neurons. TRiP offers important advantages for studying translation in a complex tissue. First, it enriches for cell type-specific translational changes, reducing the problem of heterogeneity of gene expression changes in response to a physiologically relevant stimulus. Second, TRiP improves reproducibility across samples as it is insensitive to imprecise dissection, a problem that can produce misleading results (Mathew et al. 2016).

Using TRiP we determined how a memory-inducing experience changes the translation state of CA1 pyramidal cells, a population of neurons integral to the process of memory formation. TRiP affords a nucleotide level of resolution of ribosome occupancy which allowed us to observe a striking change in ORF selection by the ribosome following animal training. By combining TRiP with animals genetically deficient in the p-eIF2α, we show that eIF2α phosphostatus significantly influences ORF selection and contributes to the ‘pro-memory’ translational state following training. These studies show that TRiP can be a powerful tool for dissecting the translational response to physiologically relevant stimuli. Furthermore, we show that TRiP has utility in determining the cell type-specific effects of translation control mutations on translation state.

### How experience alters translation state

We identified 421 ORFs that demonstrated increased translation 1h following fear conditioning training, the overwhelming majority of which were canonical proteincoding mORFs (Figure 3C). Gene ontology analysis identified components of the actin cytoskeleton among the most significantly enriched among the upregulated ORFs. Increased synthesis of proteins that are associated with the actin cytoskeleton likely supports dynamic structural plasticity observed at CA1 synapses following application of LTP protocols (Maletic-Savatic et al. 1999; Matsuzaki et al. 2004). More broadly, dynamic changes in forebrain dendritic spine structure are closely associated with a variety of forms of memory-inducing experiences (Hofer et al. 2009; Xu et al. 2009). It has been long known that neuronal activity-dependent changes in dendritic spine structure are dependent on protein synthesis (Fifková et al. 1982), but many of the actin cytoskeleton-associated proteins that are actively synthesized and promote structural plasticity have been elusive. Our results provide a wealth of candidate molecules that could play previously unidentified roles in the process of dendrite remodeling and memory formation.

The dominant class of genes demonstrating reduced translation following animal training are transcription factors. Strikingly, the IEGs Junb, Fos, Egr1, Egr4, and Npas4 all show significantly reduced translation 1h following training. These genes undergo rapid transcriptional and translational induction within minutes following a burst of neuronal activity. Rapid induction of IEGs is a desirable feature of a central regulator of memory formation, permitting rapid, activity dependent encoding of an engram or memory trace (Poo et al. 2016). However, rapid translational repression IEGs following this burst of transcriptional activity could be equally important to permit separation of memories that occur in close temporal proximity. For at least one of these transcription factors, Npas4, we have identified eIF2α dephosphorylation as a mechanism for rapid post-experience translational repression (discussed further below).

Memory-inducing experience is also known to elevate levels of bulk translation in the hippocampus (Davis and Squire 1984; Matthies 1989). The dephosphorylation of eIF2α following training that we and others (Costa-Mattioli et al. 2007) have observed may explain part of the mechanism underlying increased rates of protein synthesis. Interestingly we observed both enhanced and decreased ribosome occupancy of selected transcripts in *eIF2s1*^*S51A*/+^ mice. We suggest that one way in which eIF2α phosphostatus influences the translation of selected transcripts is through ORF selection-mediated mechanism(s). Others have proposed that gene-specific ORF selection by p-eIF2α, rather than effects on bulk protein synthesis, are important for the establishment of memory (Jiang et al. 2010). The significant overlap in differentially translated ORF resulting from training or genetic reduction in p-eIF2α suggests that the dynamic change in eIF2α phosphorylation is crucial to the ‘pro-memory’ translation state.

### eIF2α phosphorylation and ORF selection

Genetic approaches that reduce eIF2α phosphorylation lower the threshold of stimulation required for induction of LTP (Costa-Mattioli et al. 2005; 2007). Increasing p-eIF2α levels by activating the unfolded protein response (Ma et al. 2013) or through pharmacologic or pharmacogenetic means (Costa-Mattioli et al. 2007; Jiang et al. 2010) decreases the magnitude of LTP or blocks LTP completely. Consistent with changes in stimulation threshold required to achieve LTP, each of these studies show that increasing or reducing p-eIF2α either respectively impairs or enhances performance in long-term hippocampal-dependent memory tasks. Taken together, these studies show that reduced p-eIF2α levels create a translational state permissive for LTP and longterm memory formation. We observed a significant overlap between mRNAs showing significant changes in translation between trained wild-type and naïve *eIF2s1*^*S51A*/+^ mice, consistent with the concept that eIF2α dephosphorylation is permissive for memory formation. Among the mRNAs consistently demonstrating decreased translation following training and in *eIF2s1*^*S51A*/+^ mice were Htr1b and Npas4, each of which have a uORF. We demonstrate that these uORFs functionally repress translation in an eIF2α phosphorylation-dependent manner. Htr1b repression may have a functional significance in memory formation as *Htr1b^-/-^* show enhanced spatial memory (Wolff et al. 2003) and considerable evidence that HTR1B inverse agonists promote learning consolidation (Meneses 2001). Finally, treatment of rats with the HTR1B-selective antagonist SB216641 reduced latency to platform in the Morris Water Maze 28 days after training, demonstrating improved memory consolidation (Cai et al. 2013). Npas4 has a clearly established role in memory formation, including contextual fear conditioning (Sun and Lin 2016). Ablation of Npas4 in the CA3 causes severe deficits in a fear learning task while removing Npas4 from the CA1 has no effect on fear learning (Lin et al. 2008). This is consistent with eIF2α dependent-repression of Npas4 being part of the ‘pro-memory’ translation state, as the CA1-specific Npas4 knockout does not to impair establishment of fear memory. Rather, as discussed above, repression of Npas4 may play a role in discrimination of temporally close events.

The uORF/p-eIF2α regulation of mORF translation observed for Htr1b and Npas4 is likely uncommon. When observed as a whole, uORFs have little to no impact on mORF translation (Supplementary Figures 5A and 6A). However, when taken in aggregate, the presence of a dORF tends to repress the translation of the mRNA’s mORF in an eIF2α-dependent manner (Supplementary Figure 5B and 6B). While dORFs have been observed in previous studies (Calviello et al. 2015; Fields et al. 2015; Ji et al. 2015), ours is the first study to our knowledge to suggest a potential impact of dORF on mORF translation. Future studies will be required to determine if dORFs do directly impact mORF translation.

## Methods

### Animals

The *Zdhhc2*-*EGFPL10a* transgenic mouse was generated as previously described (Brichta et al. 2015). The *eIF2s1^S51A^* knock-in mice were generated and previously described by the Kaufman laboratory (Scheuner et al. 2001) and were purchased from Jackson Labs (Bar Harbor, ME). All mice were maintained on a C57BL/6 background and housed in accordance with protocols outlined by the Johns Hopkins University Animal Care and Use Committee. Animals were genotyped using primers described in the oligonucleotide table below.

### Behavioral Paradigm

The training paradigm used in this study is modeled after previous studies of hippocampal-dependent memory and translation control (Costa-Mattioli et al. 2007). Briefly, male mice 2-3 months in age were individually housed. Each mouse was handled for 5 minutes over 3 consecutive days to habituate to handling. The mice were exposed to a novel chamber for 20 minutes a day for 4 consecutive days to habituate the animal to the environment. On the eighth day of the protocol, mice were exposed to a 30 sec Tone (90 dB) that was co-terminated with a 2 sec foot shock (0.7 mA). For assessment of retention of fear memory, 24 h after training, mice were re-exposed to the environment and freezing was assessed. Mice of the appropriate genotypes were randomly assigned to trained or naïve groups using a random number table. For TRiP and puromycin release experiments, following the training period mice were returned to their home cage for 1 h and then sacrificed by decapitation. Hippocampi were rapidly dissected in buffer containing cycloheximide (Kulicke et al. 2014) and then snap-frozen on ethanol-dry ice bath and stored at −80°C for downstream studies. For puromycin release samples, cycloheximide was omitted from the dissection buffer.

### Quantitative reverse transcription PCR

Reverse transcription (RT) was performed on 100 ng of TRAP purified and input RNA using Superscript III (Thermofisher) and random hexamers using the manufacturer’s protocol. RT reaction was diluted 1:10 with water and 0.5 μl of cDNA was used in each real-time PCR assay. Taqman predesigned assays were used for *Camk2a*, *Gad2*, and *Gfap* using Applied Biosystems Taqman Master Mix (Thermofisher). *Egfp* and *Rps2* assays used custom primers and were applied using Sybr green master mix (Thermofisher). Primers for these assays are summarized in the oligonucleotide table. All assays were performed in quadruplicate from 3 sets of TRAP purifications and their inputs (n = 3) on a Viia7 real-time PCR system (Thermofisher).

### Translating Ribosome Affinity Purification and Ribosome Profiling (TRiP)

Hippocampi of two *Zdhhc2*-*EGFPL10a* or *Zdhhc2*-*EGFPL10a; eIF2s1*^*S51A*/+^ mice were pooled and homogenized in a pre-chilled Dounce homogenizer. TRAP was performed as previously described using all the suppliers of reagents specified in the previous description (Kulicke et al. 2014). Following final high-salt wash, beads bound with tagged ribosomes were resuspended in 250 μl low salt wash (10 mM HEPES, 150 mM KCl, 5 mM MgCl_2_, 1% NP-40). Following resuspension, 7 μl RNase I (Ambion, Thermofisher) was added and RNA digested for 45 minutes at 24°C with agitation (700 RPM, Eppendorf Thermomixer). RNase digestion was terminated by addition of 750 μl of Trizol LS (Thermofisher) to the bead slurry. RNA was isolated following the manufacturer’s protocol.

To aid in the efficient recovery of ribosome protected fragments, 100 pmol of RNA oligos (EM6/7, see oligonucleotide table below) bearing an I-S*ce*I homing endonuclease site were added to each sample. Ribosome profiling libraries were generated as previously described (Ingolia et al. 2012) with the following two modifications. First, following recovery of ribosome protected fragments and prior to dephosphorylation and adapter ligation, rRNA subtraction was performed using the NEBNext rRNA depletion kit (NEB). Second, after PCR amplification, libraries were digested if I-S*ce*I (NEB) to remove carrier RNA sequences. After digest, full length libraries were recovered by gel extraction as described (Ingolia et al. 2012). Library quality was assessed by BioAnalyzer (Agilent) and quantitated by qPCR (KAPA Biosciences). Libraries were sequenced on a HiSeq 2000 (Illumina) at the Lieber Institute and the Carnegie Institute Department of Embryology.

### Bioinformatics

*Read processing and mapping:* Adapter sequences were trimmed from reads using FASTX trimmer (http://hannonlab.cshl.edu/fastx_toolkit/). Ribosomal RNA (rRNA) sequences were removed using Bowtie2 (Langmead and Salzberg 2012) and the resulting FASTQ file was passed to Tophat2 (Kim et al. 2013) for alignment to a preassembled transcriptome. For preliminary experiments (Fig 1) reads were aligned to the mm10 assembly and the gene models retrieved knownGenes table from the UCSC genome browser. For all other mapping (Figs 2-5), FASTQ files were mapped to Gencode release M9 transcriptome, using the APPRIS_Principal transcript annotation. Where more than one APPRIS_Principal transcript was identified, the longest transcript was selected.

*ORF annotation:* The mapped reads from all TRiP samples were combined into a single BAM file and analyzed using Ribotaper (Calviello et al. 2015). Ribosome protected fragments 28-30 nt were well phased based on metagene analysis of annotated start and stop codons and were subsequently used to identify translated segments of the transcriptome. Output of Ribotaper was reanalyzed to identify overlapping uORFs, a feature not currently available in the release used (v1.3). Overlapping uORFs were identified as ORFs overlapping the canonical ORF but in a different reading frame. ORFs demonstrating a Ribotaper *p* value <0.05 were included in subsequent analysis to annotate ORF translation.

*ORF assignment and differential gene expression analysis:* Ribosome protect fragments were assigned to Ribotaper-identified ORFs using a custom R script. The per-ORF read count table was passed to edgeR (v3.1.4) for differential expression analysis using the generalized linear model function (Robinson et al. 2010). Dimension reduction analysis was performed using the `plotMDS` function within edgeR and `prcomp` function within the R stats package.

### Western blot

Total lysate were prepared from mouse hippocampus by homogenization in RIPA buffer supplemented with complete EDTA-free protease inhibitors (Roche) and phosphatase inhibitors (cocktails 2 and 3, Sigma). Lysates were quantitated by BCA assay (Thermofisher) and boiled in SDS-PAGE sample buffer. Samples (20 μg per lane) were run on tris-glycine polyacrylamide gels and transferred to PVDF (Bio-Rad). Membranes were blocked in 5% non-fat dry milk or 3% BSA in 1x PBS+ 0.1% Tween-20. Western blotting for puromycin release assay was performed from the lysates described in that assay. See immunoreagent summary table for details of antibodies used.

### Puromycin release assay

This variation on the SUnSET method (Schmidt et al. 2009) is adapted from a previously described protocol (Biever et al. 2015). In brief, frozen hippocampus is homogenized in 400 μl of puromycin release buffer (10 mM HEPES pH 7.4, 5 mM MgCl_2_, 150 mM KCl, 200 μM emetine, 0.5 mM DTT, EDTA-free protease inhibitors, 200 μM ATP, 100 μM GTP) in a motorized Dounce homogenizer. The lysate is clarified by spinning at 12,000 x g at 4°C for 10 minutes. 300 μl of clarified lysate is transferred to a new tube on ice and supplemented with puromycin to final concentration of 1.25 mM. Following 10 min incubation on ice, the reaction is terminated by the addition of 15 μl of 500 mM EDTA.

### Primary Mouse Embryonic Fibroblast Preparation

Embryos were harvested at 15 days post coitum from *eIF2s1*^*S51A*/+^ females bred with an *eIF2s1*^*S51A*/+^ male. Individual embryos were decapitated and the visceral organs removed prior to enzymatic digestion, reserving some tissue for genotyping. Cells recovered following digestion were plated in a T25 flask in standard media (DMEM+10% FBS, pen-strep, Thermofisher/Gibco). Cultures were expanded following identification of *eIF2s1^S51A/S51A^* homozygotes and wild-type littermates. Cells were passaged for up to 4 times before discarding.

### Luciferase Reporter Assay

Luciferase reporters were generated by inserting the 5’-UTR sequences of *Htr1b* or *Npas4* upstream of the *Renilla* luciferase (Rluc) ORF on the pRL-TK plasmid (Promega). Actb 5’-UTR reporter was obtained from Addgene(Thoreen et al. 2012). To ensure accurate transcription initiation, the 5’-UTR-Rluc fusion was amplified with primers adding a T7 RNA polymerase reporter and a poly(A_60_) to the 3’-UTR sequence. These PCR products were used as a template for *in vitro* transcription and capping of Rluc mRNA (HiScribe T7 ARCA mRNA kit, NEB). The same strategy was used to generate a firefly luciferase (Flu) mRNA reporter with a generic 5’-UTR sequence (pGL4.53, Promega). MEFs were plated at a density of 50,000 cells/well in a poly-L-ornithine-coated 24-well plate. After 24 h, cells were transfected with 200 ng of Rluc and 200 ng of Fluc reporter mRNA (Lipofectamine 2000, Thermofisher). Cells were harvested in 1×PLB (Promega) 16 h following transfection. Luciferase activity was measured using Dual Luciferase kit (Promega). Luciferase assays were performed using MEFs from three different embryos and 2 or 3 independent transfections (n = 6 to 9 per group).

### Statistical analysis

Statistical tests were performed either using Prism (v 5, Graphpad) or R. Statistical assessment of TRiP data was made using read counts data and GLM multiple comparison testing in edgeR.

## Acknowledgements

We thank Eric Mills for helpful discussion concerning low-input ribosome profiling approaches and Lorenzo Calviello for assistance in running Ribotaper. We also thank Jungwoo Wren Kim for thoughtful discussion and comments on the manuscript. Illustrations in Figures 1A, 3A, and S3 were made by I-Hsun Wu. This work was supported by NIDA grant DA00266. SME, TMD, and VLD acknowledge the joint participation by the Diana Helis Henry Medical Research Foundation Parkinson’s Disease Program through its direct engagement in the continuous active conduct of medical research in conjunction with the Johns Hopkins Hospital and the Johns Hopkins University School of Medicine and the Foundation’s Parkinson’s Disease Program H-2015. TMD and PG are supported by the JPB Foundation. KC is supported by a NINDS Diversity supplement for P50NS038377. TMD. is the Leonard and Madlyn Abramson Professor in Neurodegenerative Diseases.

## Competing Interests

The authors have no competing interests to declare.

Immunoreagents.

Immunoreagents
| Antibody | Source | Catalog # | Use |
| --- | --- | --- | --- |
| Mouse anti-EGFP | Memorial Sloan Kettering Monoclonal Antibody Facility | HtzGFP-19C8 | TRAP (50 µg/ IP) |
| Mouse anti-EGFP | Memorial Sloan Kettering Monoclonal Antibody Facility | HtzGFP-19F7 | TRAP (50µg/ IP) |
| Goat anti-EGFP | Rockland Antibodies | 600-109-215 | Immunofluorescence (1:1000) |
| Rabbit anti-p-eIF2α | Cell Signaling Technologies | D9G8 | Western blot (1:1000) |
| Rabbit anti-eIF2α | Cell Signaling Technologies | D7B3 | Western blot (1:2000) |
| Mouse anti-Puromycin | EMD Millipore | MABE343 | Western blot (1:10,000) |
| Mouse anti-Tubulin | Sigma | T8328 | Western blot (1:10,000) |

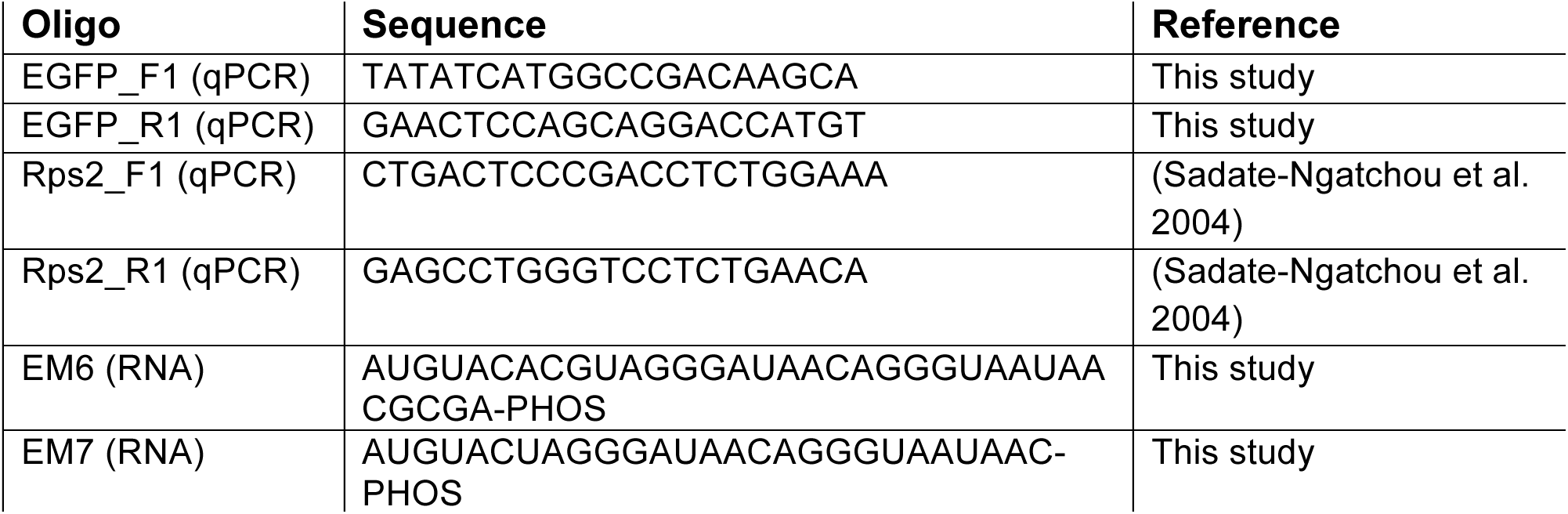
Oligonucleotides.

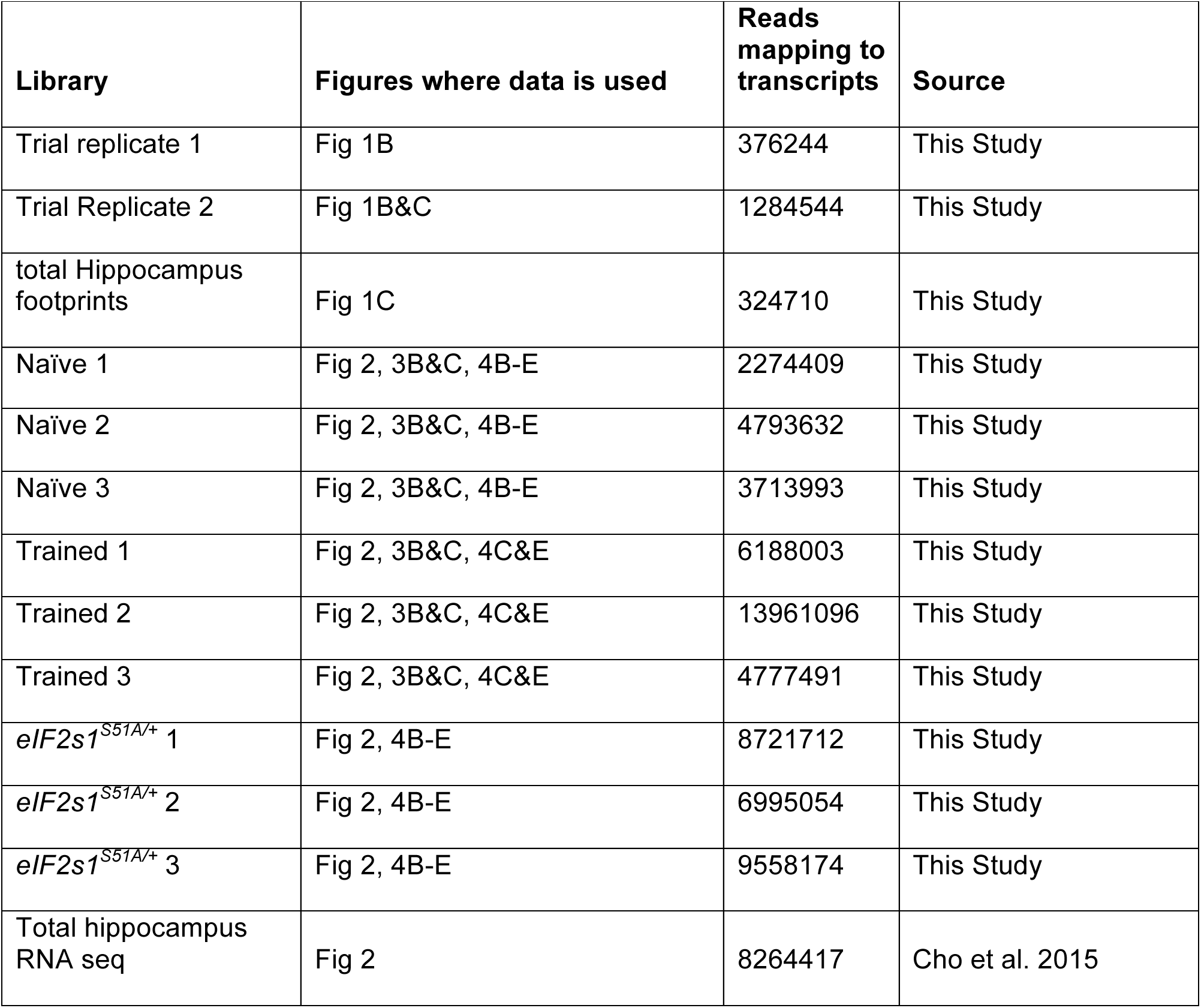
Datasets used in this study.

---

**Supplementary Figure 1:**
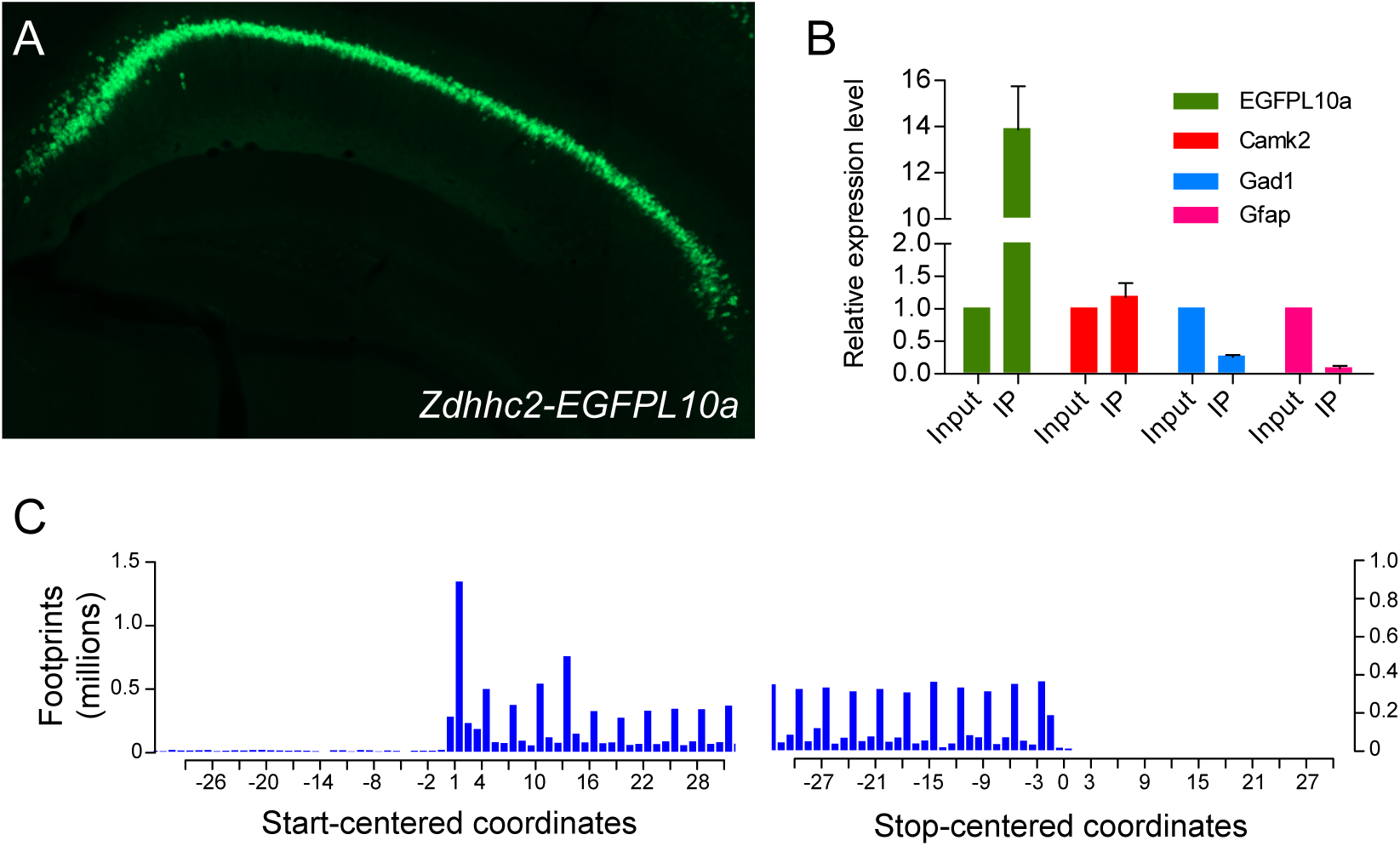
Characterization of *Zdhhc2*-*EGFPL10a* transgenic mouse. (A) Anti-EGFP immunofluorescence shows prominent expression in pyramidal cells of the CA1. (B) TRAP qPCR testing enrichment/ depletion of marker genes comparing immunopurified mRNA to input (n = 3). (C) Representative metagene analysis of TRiP ribosome-protected fragments.

**Supplementary Figure 2:**
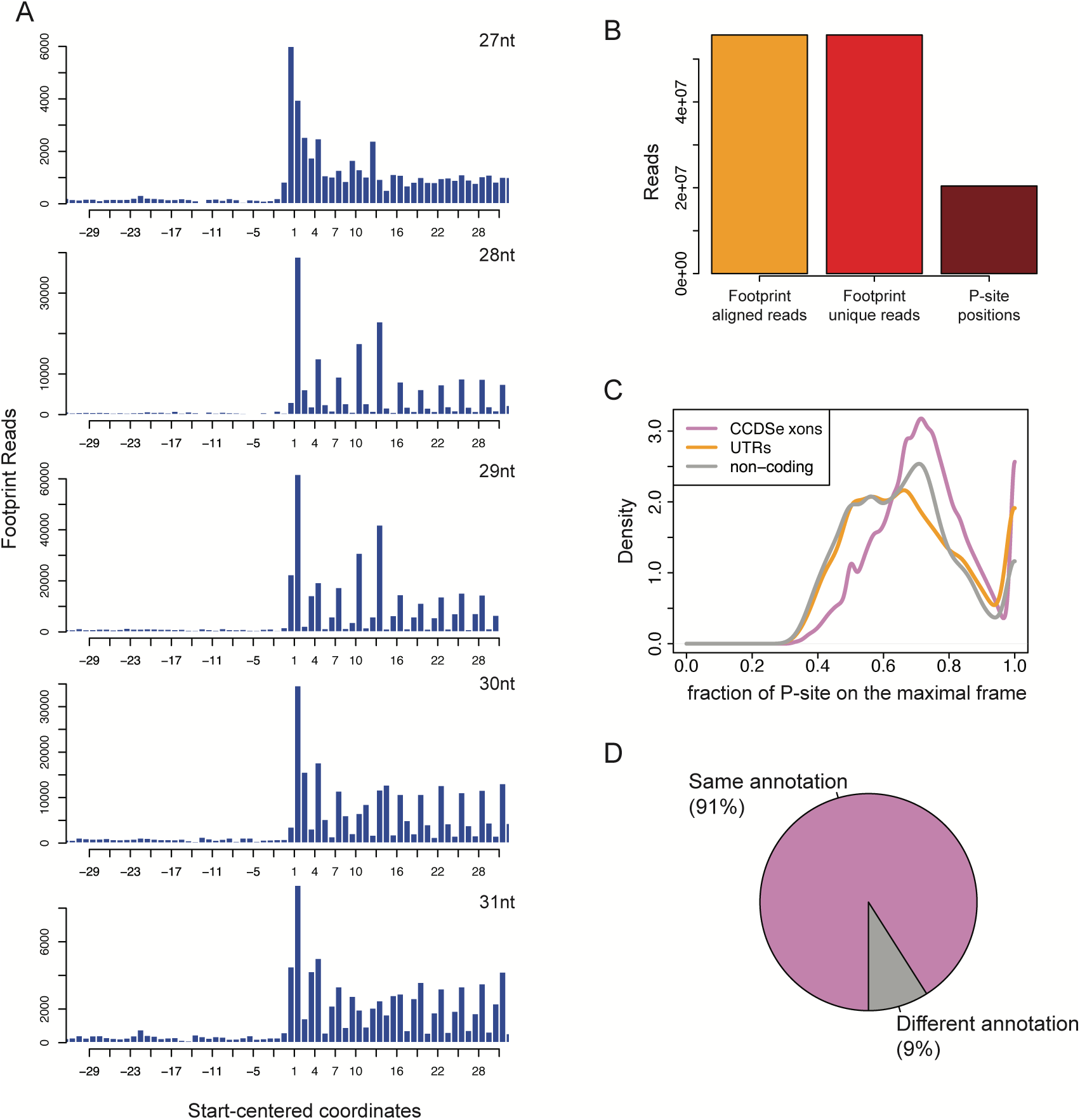
Quality checks from Ribotaper analysis of TRiP data. (A) Metagene analysis of ribosome protected fragments of various lengths shows the highest periodicity are of fragments 28-30 nt in length. (B) Summary of footprints analyzed and used to predict actively translated ORFs. (C) Enrichment of in-frame footprints in CCDS exons compared with ncRNA or UTRs. (D) Agreement between Ribotaper-predicted and CCDS annotated ORFs.

**Supplementary Figure 3:**
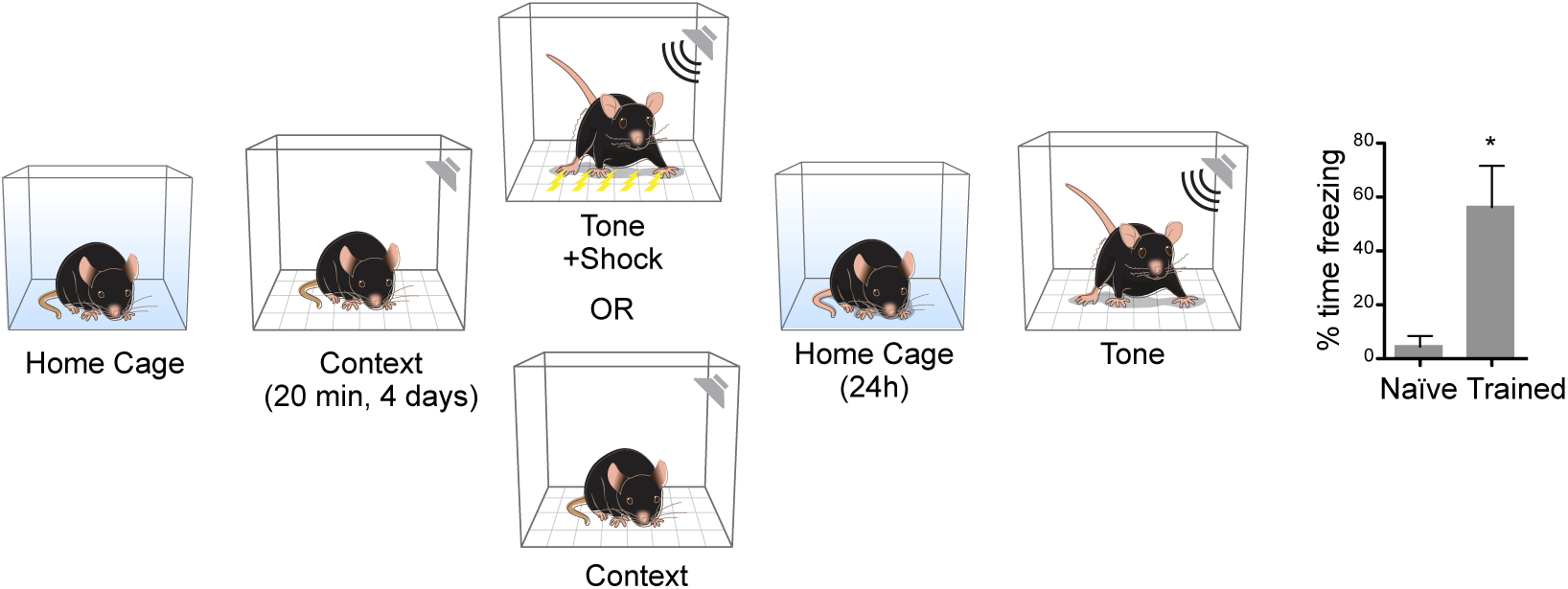
Fear training instills long-term fear memory. Animals were handled as described in methods. Following day 4 of habituation, animals were either exposed to the context (naïve group) or exposed to context, provided a 30 sec tone co-terminating with a 2” shock. Mice were returned to home cage for 24h and memory was assessed by percent time freezing (*p* <0.05, n= 4 per group). ^∗^ *p* < 0.05, Paired T-test.

**Supplementary Figure 4:**
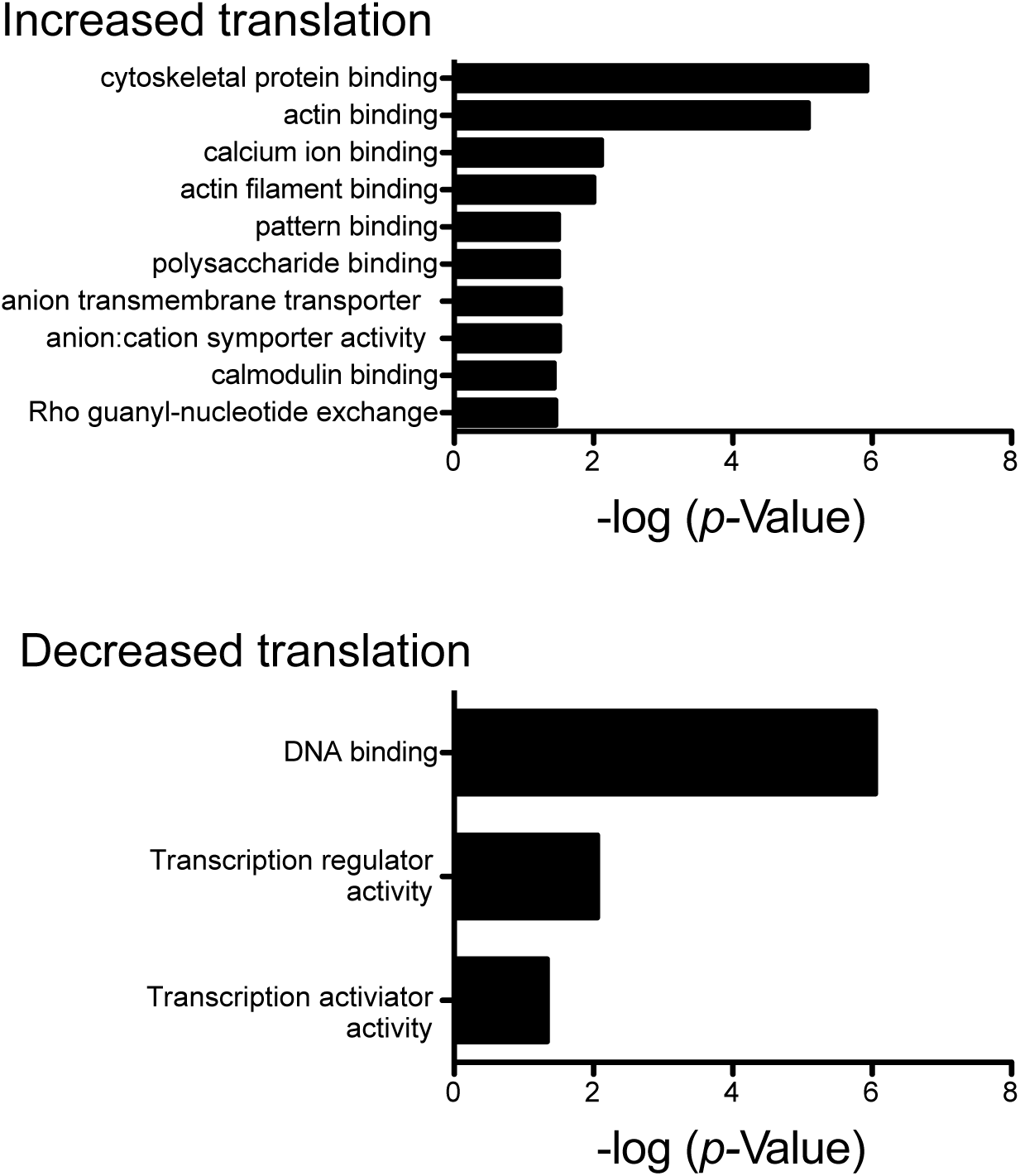
Gene ontology analysis of mRNAs showing differential translation following training. The molecular function of genes showing statistically significant changes in translation following training were analyzed using DAVID (v 6.7) (Huang et al. 2008). Significant enrichment of molecular functions was assess at a Benjamini-correct *p* value of <0.01.

**Supplementary Figure 5:**
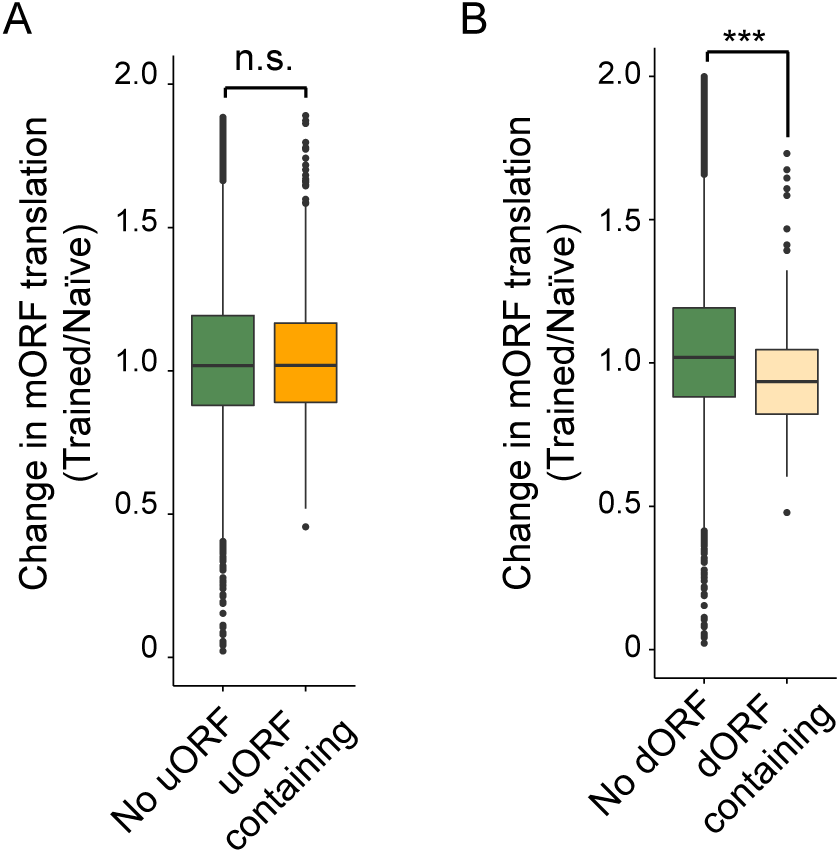
Influence of uORFs and dORFs on mORF translation following training. (A) The change in translation of mORFs whose mRNA does or does not contain a uORF. (B) The change in translation of mORFs whose mRNA does or does not contain a dORF (*p* < 6.4 × 10-9, Wilcoxon rank sum tests). Center bar indicates median fold-change and box indicates interquartile range.

**Supplementary Figure 6:**
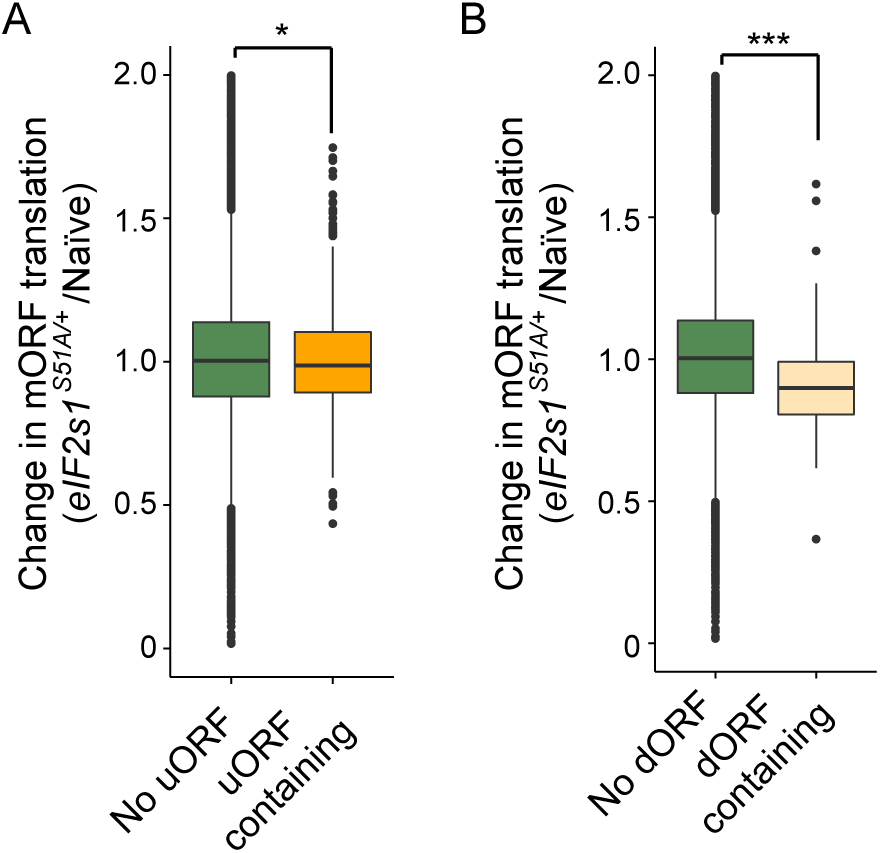
Influence of uORFs and dORFs on mORF translation due to genetic reduction of eIF2α phosphorylation. (A) The change in translation of mORFs whose mRNA does or does not contain a uORF(^∗^, *p* < 0.05, Wilcoxon rank sum tests). (B) The change in translation of mORFs whose mRNA does or does not contain a dORF (^∗∗∗^, *p* < 6.4 × 10-9, Wilcoxon rank sum tests). Center bar indicates median fold-change and box indicates interquartile range.

**Supplementary Figure 7:**
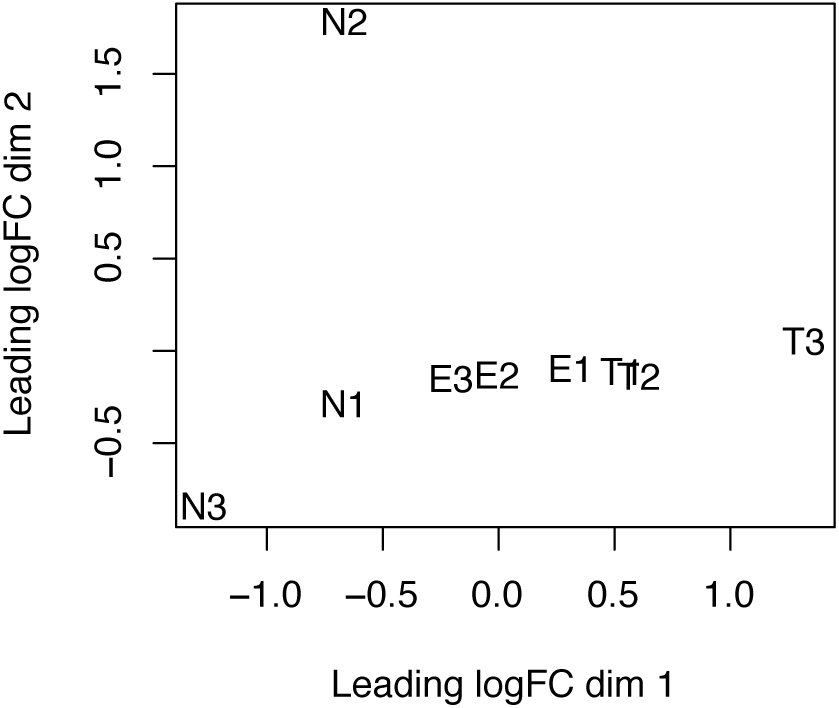
Multidimensional scaling plot comparing profiles of naïve, trained, and *eIF2s1S51A*/+ TRiP data. Dimensional reduction based leading logfold change in ORF-specific translation across replicates of naïve (N1-3), trained (T1-3), and naïve *eIF2s1S51A*/+ (E1-3) TRiP experiments.

**Supplementary Figure 8:**
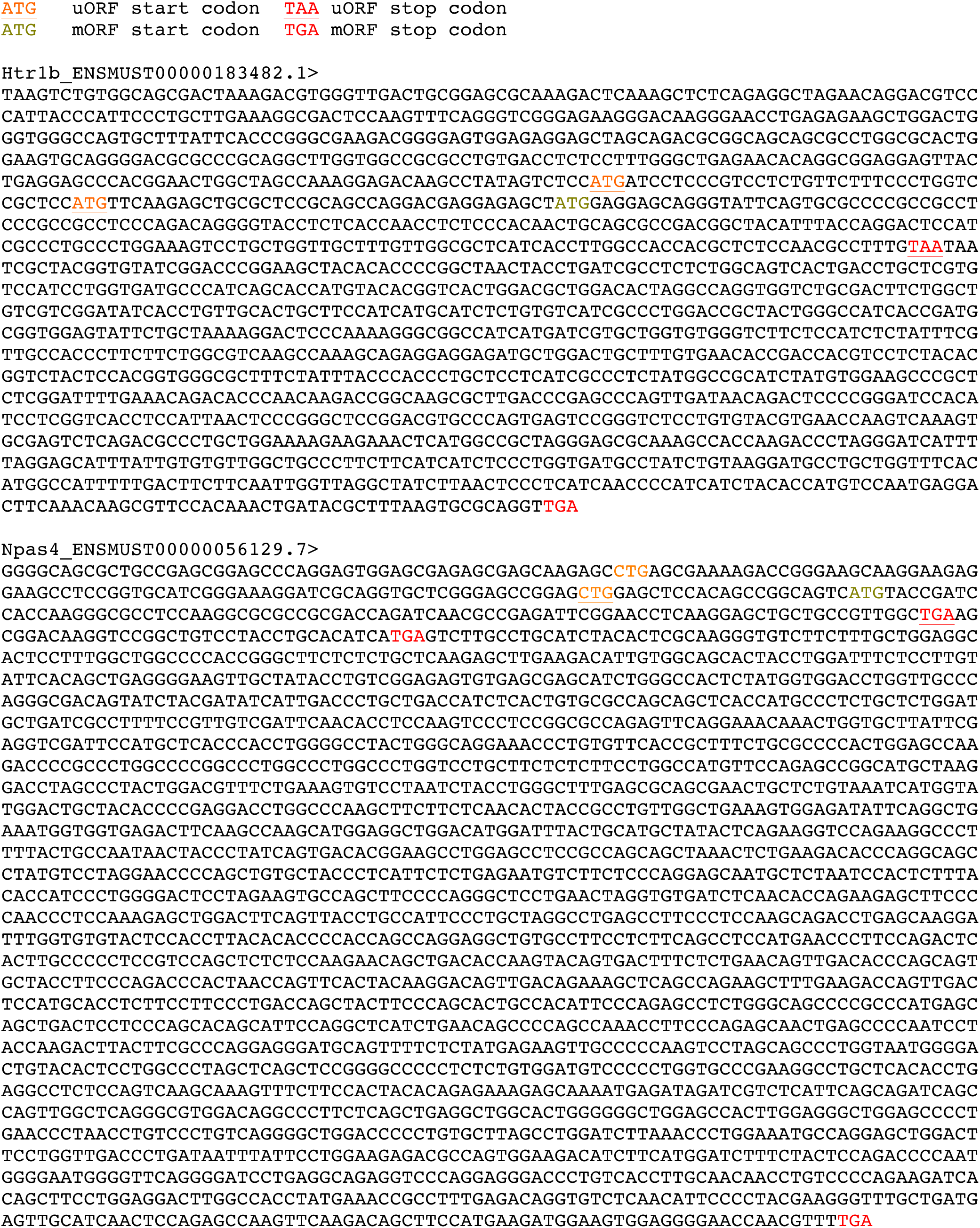
5’UTR and mORF sequence of Htr1b and Npas4. Start and stop codons of uORFs are underlined. Start codons of mORF are in green and stop codons in red.

